## Supporting Information Appendix for "Interleukin 10 controls the balance between tolerance, pathogen elimination and immunopathology in birds"

**A**

```
ATG CAG ACC TGC TGC CAA GCC CTG TTG CTG CTG CTG GCT GCA TGC ACC CTG CCT GCC CAC
 M   Q   T   C   C   Q   A   L   L   L   L   L   A   A   C   T   L   P   A   H

TGC TTG GAG CCC ACC TGC CTG CAC TTC TCT GAG CTG CTG CCC GCC CGG CTG CGG GAG CTG
 C   L   E   P   T   C   L   H   F   S   E   L   L   P   A   R   L   R   E   L

AGG GTG AAG TTT GAG GAA ATT AAG GAC TAT TTT CAA TCC AGA GAC GAT GAA CTT AAC ATC
 R   V   K   F   E   E   I   K   D   Y   F   Q   S   R   D   D   E   L   N   I

CAA CTG CTC AGC TCT GAA CTG CTG GAT GAG TTT AAG GGG ACC TTT GGC TGC CAG TCT GTG
 Q   L   L   S   S   E   L   L   D   E   F   K   G   T   F   G   C   Q   S   V

TCA GAG ATG CTG CGC TTC TAC ACA GAT GAG GTC CTG CCC CGT GCC ATG CAG ACC AGC ACC
 S   E   M   L   R   F   Y   T   D   E   V   L   P   R   A   M   Q   T   S   T

AGT CAT CAG CAG AGC ATG GGC GAC CTG GGC AAC ATG CTG CTG GGC CTG AAG GCG ACG ATG
 S   H   Q   Q   S   M   G   D   L   G   N   M   L   L   G   L   K   A   T   M

CGG CGC TGT CAC CGC TTC TTC ACC TGC GAG AAG AGG AGC AAA GCC ATC AAG CAG ATC AAG
 R   R   C   H   R   F   F   T   C   E   K   R   S   K   A   I   K   Q   I   K

GAG ACG TTC GAG AAG ATG GAT GAG AAC GGG ATC TAC AAA GCC ATG GGG GAG TTC GAC ATC
 E   T   F   E   K   M   D   E   N   G   I   Y   K   A   M   G   E   F   D   I

TTC ATC AAC TAC ATC GAG GAG TAC CTG CTG ATG AGG AGG AGG AAG TGA
 F   I   N   Y   I   E   E   Y   L   L   M   R   R   R   K   *
```

**B**

```
        ACC TGC TGC CAA GCC CTG TTG CTG CTG CTG GCT GCA TGC ACC CTG CCT GCC CAC
ATG CAG ACC TGC TGC CAA GCC CTG TTG CTG CTG CTG GCT GCA TGC ACC CTG CCT GCC CAC
 M   Q   T   C   C   Q   A   L   L   L   L   L   A   A   C   T   L   P   A   H

                      C CTG CAC TTC TCT GAG CTG CTG C
TGC TTG GAG CCC ACC Tag gTG CAC TTC TCT GAG CTG CTG CCC GCC CGG CTG CGG GAG CTG
TGC TTG GAG CCC ACC TGC CTG CAC TTC TCT GAG CTG CTG CCC GCC CGG CTG CGG GAG CTG
 C   L   E   P   T   C   L   H   F   S   E   L   L   P   A   R   L   R   E   L

AGG GTG AAG TTT GAG GAA ATT AAG GAC TA
AGG GTG AAG TTT GAG GAA ATT AAG GAC TAT TTT
 R   V   K   F   E   E   I   K   D   Y   F
```

**C**

```
ATG CAG ACC TGC TGC CAA GCC CTG TTG CTG CTG CTG GCT GCA TGC ACC CTG CCT GCC CAC
 M   Q   T   C   C   Q   A   L   L   L   L   L   A   A   C   T   L   P   A   H

TGC TTG GAG CCC ACC TAG
 C   L   E   P   T   *
```

**Figure S1: Wildtype and edited IL10 sequences for IL10KO edit. A**: Full-length chicken IL10 protein sequence (blue; 175 amino acids; NP_001004414.2) aligned to *IL10* coding sequence. Alternating exons are shown in black and green; the start codon is underlined; the stop codon is shown in red (*). **B**: Chicken *IL10* exon 1 sequence (black), showing the sequence and position of gRNA ggIL10_exon1_g3 (green box), Protospacer Adjacent Motif (PAM; yellow box), and ssODN IL10_exon1_HDRoligo2 (grey box), with the three substituted nucleotides (red, lowercase). **C**: Edited chicken *IL10* exon 1 showing the premature stop codon in red (*) and truncated IL10 peptide (25 amino acids).

```
                  IL10_enhancer_guide1
                  GTTTCGTAGCGGGTGAATGAAGG
WT      AATGGGTGTTTCGTAGCGGGTGAATGAAGGGCAGCAAAGTCTCCACTTCCCCGGTGTCACTGACAGCAAA
Edited  AATGGGTGTTTCGTAGCGGGTGA----------------533 bp deletion----------------

WT      AAGGAATGGATGAGAAGTGGGAAATTTGGGGTAAGAAACATACTGTTAAGCAGCCTGGAAATCTAGGCAA
Edited  ---------------------------533 bp deletion----------------------------

WT      TTCCTGCAGCTGTATCTATAATAGTGCATAAAATACACCGTGTTCCCAAAGGCAATGATTGAGACCCAGA
Edited  ---------------------------533 bp deletion----------------------------

WT      GGGGGAAAAATGGGAGAAGTTGCCCCTATGGAAGCTGCCGGCATCAAAGAGTGCAAAAGGCTTTTGAGTA
Edited  ---------------------------533 bp deletion----------------------------

WT      ATGTTTCCTTCCTGAAATGATCTTCTTGGCAATCATGAAAGCTGTTTCCTAAAGGTTTAGAAATACACCC
Edited  ---------------------------533 bp deletion----------------------------

WT      ACGTCCTCGGACACTCTTTAGGAAAGGTGCTTTATTCATTCATGCAACTGCTTCGTAGTACTGCAGAGTC
Edited  ---------------------------533 bp deletion----------------------------

                                       AP1
WT      AATGCGTTTCCACAGCAGTCAGCAGTGGTGAGTCAGAGCGAAACTGCACCGTCAAAATCTCTGCCATGGT
Edited  ---------------------------533 bp deletion----------------------------

                                                  IL10_enhancer_guide2
                     FOXP3                        GTGCAGGGCAGTTTCCTTTGTGG
WT      GAAGGAAACGCAGGTAAACACAGCGAGCATCGTGCGTGTGCAGGGCAGTTTCCTTTGTGGAATGGACACG
Edited  ---------------------------533 bp deletion-----------------------CACG

WT      ACTGGCTTACATGGTGTGCAGCTGAAAAACAAAGCAGGTTCCAAAACAGGGAGGAGGGAACTGCATAAGG
Edited  ACTGGCTTACATGGTGTGCAGCTGAAAAACAAAGCAGGTTCCAAAACAGGGAGGAGGGAACTGCATAAGG
```

**Figure S2: Wildtype and edited *IL10* putative enhancer sequences for IL10EnKO edit.** The chicken *IL10* putative enhancer region (548 bp) is shown in blue, with the core conserved region (SI Appendix, Figure S3) double-underlined. The turquoise and pink boxes highlight the location of the AP1 and FOXP3 sites, respectively. The position and sequence of gRNAs IL10_enhancer_guide1 and IL10_enhancer_guide2 and corresponding PAMs are indicated by green and yellow boxes, respectively. The 533-bp sequence deleted in IL10EnKO PGCs is indicated by a grey box.

```
Chicken       TGAAATGATCTTCTTGGCAATCATGAAAGCTGTTTCCTAAAGG-TTTAGAAATACACCCACGTCCTC-GGACACTCTTTA
Budgerigar    TGAAATGATCTGCTTGGCAAGCATGAAAGCTGTTTCCTAAAGGGCTCAGAAATACACCCACATCCCCCGGACACTCTTTA
Buzzard       TGAAACGATCTTCTTGGCAAACATGAAAGTTGCTTCCTAAAGG-TTTAGAAATACACCCACGTCCCCCGGACACTCTTTA
Canary        TACAGCGATCTTCTTGGCGAAGCCGAGAGCCGTTTCCTAAAGG-TTCGGCGCTACACCCACGGCCCCCGGACACTCTTTA
Duck          TGAAACGATCTTCTTGGCAACTGTGAAAGCTGTTTCCTAAAGG-TTTAGAAATACACCCACGTCCCC-GGACACTCTTTA
Eagle         TGAAACAATCTTCTTGGCAAACATGAAAGCTGCTTCCTAAAGG-TTTAGAAATACACCCACGTCCCCCGGACACTCTTTA
Emu           TGAAATGATCTTCTTGGCAAACATGAAAGCTGTTTCCTAAAGG-TTTAGAAATACGCCCACGTCCCC-GGACACTCTTTA
Kestrel       TGAAATGATCTTCTTGGCAAACATGAAAGCTGTTTCCTAAAGG-TTTAGAAATACACCCACATCCCCCGGACACTCTTTA
Kiwi          TGAAATGATCTTCTTGGCAAACATGAAAGCTGTTTCCTAAAGG-TTTAGAAATACACCCACATCCCC-GGACACTCTTTA
Ostrich       TGAAATGATCATCTTGGCAAATATGAAAGCTGTTTCCTAAAGG-TTTAGAAATACACCCACGTCCCC-GGACACTCTTTA
Pheasant      TGAAATGATCTTCTTGGCAATCATGAAAGCTGTTTCCTAAAGG-TTTAGATATACACCCACGTCCTC-GGACACTCTTTA
Ruff          TGAAATGATCTTCTTGGCAAACGT-AAAGCTGTTTCCTAAAGG-TTTAGAAATACACCCACATCCCCCGGACACTCTTTA
Sandpiper     TGAAATGATCTTCTTGGCAAACGT-AAAGCTGTTTCCTAAAGG-TTTAGAAATACACCCACATCCCCCGGACACTCTTTA
Turkey        TGAAATGTTCTTCTTGGCAATCATGAAAGCTGTTTCCTAAAGG-TTTAGAAATACACCCACGTCCTC-GGACACTCTTTA
Zebrafinch    TGAAGCGATCTTCTTGGCAAAGACAAAGCTGTTTCCTAAAGG-CTCAGCGCTACACCCACGGCCCCCGGACACTCTTTA
              *  *    **  ****** *     * **  * **********  *  *   *** *****  ** * ************

Chicken       GGAAA-GGTGCTTTATTCATTCATGCAACTGCTTCGTAGTACT-GCAGAGTCAATGCGTTTCCACAGCAGTCAGCAGTGG
Budgerigar    GGAAAAGGTGCTTTATTCATTCATGCAACTGCTTTGTAAGGCT-GCGGAGTCAATGCATTTCCACAGCAGTAAGCACAAG
Buzzard       GGAAAAGGTGCTTTATTCATTCATGCAACCACTTCGTAGTACT-GCAGAGTCAATGCATTTCCACAGCAGTAAGCAAGAG
Canary        GGAAAAGGTGTTTGCTTCATTCATGAAGCT-CTCTGTA--GTA-GTGCAGCCACCGCCTTTCCACAGCGCTCAGCAGGGC
Duck          GGAAAAGGTGCTTTATTCATTCATGCAAGCACTTCGTAGTACT-GCAGAGTCAATGCATTTCCACAGCAGTAAGCAAGGG
Eagle         GGAAAAGGTGCTTTATTCATTCATGCAACCACTTCGTAGTACT-GCAGAGTCAATGCATTTCCACAGCAGTAAGCAAGAG
Emu           GGAAAAGGTGCTTTATTCAYTCATGCAACCACTTCGTAGTACT-GCAGAGTGAATGTATTTCCACAGCAGTAAGCAAGAG
Kestrel       GGAAAAGGTGCTTTATTCATTCATGCAACCACTCCGTAGTACTTGCAGAGTCAATGCCTTTCCACAGCAGTAAGCAAGAA
Kiwi          GGAAA-GGTGCTTTATTCATTCATGCAACCACTTCGTAGTACT-GCAGAGTGAATGTATTTCCACAGCAGTAAGCAAGAG
Ostrich       GGAAAAGGTGCTTTATTCATTCATGCAACCACTTCGTAGTACT-GCAGAGTGATTGCATTTCCACAGCAGTAAGCAAGAC
Pheasant      GGAAA-GGTGCTTTATTCATTCATGCAACAGCTTCGTAGTACT-GCAGAGTCAATGCGTTTCCACAGCAGTAAGCAGTGG
Ruff          GGAAAAGGTGCTTTATTCATTCATGCAACTGCTTCGTAGTACT-GCAGAGTCAATGCCTTTCCACGGCAGTAAGCAGGGG
Sandpiper     GGAAAAGGTGCTTTATTCATTCATGCAACCGCTTCGTAGTACT-GCAGAGTCAATGCCTTTCCACGGCAGTAAGCAGGAG
Turkey        GGAAA-GGTGCTTTATTCATTCATGCAACTACTTCGTAGTACT-GCAGAGTCAATGTGTTTCCACAGCAGTAAGCAGTGG
Zebrafinch    GGAAAAGGTGTTTTCTTCATTCATGAATCT-TGCTGTA--GTA-GAGCAGCCACCGCCTTTCCACAGCGCTCAGCAGGGC
              ***** **** **  **** ***** *        ***      *   **  *  *  ******* **  * ****
              AP1                                                    FOXP3
Chicken       TGAGTCAGAGCGAAACTGCACCGTCAAAATCTCTGCCATGGTGAAGGAAACGCAGGTAAACACAGC-GAGCATCGTGCGT
Budgerigar    TGAGTCAGAGCGAAACTGCACCGTCAAGATCTCTGACATGGCAAAGGAAACCCAGGTAAACACAGCAGAGTATCGGTCCT
Buzzard       TGAGTCAGAGCGAAACTGCACTGTCAAGATCTCTGACACGGCAAAGGAAATACAGGTAAACACAGCAGAGTATCATCCGT
Canary        TGAGTCACAGCGAAACTGCACCGTCAAGATCCCTGACGAGGCAGAGGAAACGCAGGTAAACACAGCAGAGAGTCG-CCCT
Duck          TGAGTCAGAGCGAAACTGCACCGTCAAGATCTCTGACACGGCGAAGGAAACGCAGGTAAACACAGCCGAGTGCCGTGCGT
Eagle         TGAGTCAGAGCGAAACTGCACTGTCAAGATCTCTGACACGGCAAAGGAAATACAGGTAAACACAGCAGAGTATCATCCGT
Emu           TGAGTCAGAGTGAAACTGCACCGTCAAGATCTCTGACATGGCAAAGGAAATACAGGTAAACACAGTATAGTATCATCAGT
Kestrel       TGAGTCAGAGTGAAACTGCACCGTCAAGATCTCTGACACGGCAAAGGAAATACAGGTAAACACAGCAGAGTAGCATCCGT
Kiwi          TGAGTCAGAGCGAAACTGCACYGTCAAGATCTCTGACACGGCAAAGGAAATACAGGTAAACACAGTACAGTATCATCAGT
Ostrich       TGAGTCAGAGTGAAACTGCACCGTCAAGATCTCTGACATGGCAAAGGAAATACAGGTAAACACAGTACAGTATCATCAGT
Pheasant      TGAGTCAGAGCGAAACTGCACCGTCAAAATCTCTGCCATGGCGAAGGAAACACAGGTAAACACAGC-GAGCATCGTGCGT
Ruff          TGAGTCAGAGCGAAACTGCACCGTCAAGATCTCTGACACGGCGAAGGAAACACAGGTAAACACAGCGGGGTGGCATCCGT
Sandpiper     TGAGTCAGAGCGAAACTGCACCGTCAAGATCTCTGACACGGCGAAGGAAATACAGGTAAACACAGCGGGGTGGCATCCGT
Turkey        TGAGTCAGAACGAAACTGCACCGTCAAAATCTCTGCCATGGCGAAGGAAACACAGGTAAACACAGC-GAGCATCGTGCGT
Zebrafinch    TGAGTCACAGCGAAACTGCACCGTCAAGATCTGTGACGGGGCAGAGGAAACGCAGGTAAACACAGCAGAGAATCG-CCCT
              ******* *  ********** ***** ***  ** *  **   ******  *************    *   *     *

Chicken       GTGCAGGGCA--GTTTCCTTTGTGGAATGGACACGACT
Budgerigar    GCGCAGGCCAGAGTTTCCTTTGTGGAATGGACACATCT
Buzzard       GCGCAGGCCAGAGTTTCCTTTGTGGAATGGACACATCT
Canary        GCGCACGCCAGAGTTTCCTCTGCGGAACGGGCACGGCC
Duck          GCGCAGGACG--GTTTCCTTTGTGGAATGGACACATCT
Eagle         GCGCAGGCCAGAGTTTCCTTTGTGGAATGGACACATCT
Emu           GCGCAGGCCAGAGTTTCCTTTATGGAACGGACACATCT
Kestrel       GCGCAGGCTAGAGTTTCCTTTGTGGAATGGACACATCT
Kiwi          GCGCAGGCCAGAGTTTCCTTTATGGAATGGACACATCT
Ostrich       GCGCAGGCCAGAGTTTCCTTTATGGAATGGACACATCT
Pheasant      GTGCAGGGCA--GTTTCCTTTGTGGAATGGACGCAACT
Ruff          GCGCAGGCCGCAGTTTCCTTTGTGGAATGGACACATGG
Sandpiper     GCGCAGGCCACAGTTTCCTTTGTGGAATGGACACATCG
Turkey        GTGCAGGGCA--GTTTCCTTTGTGGAATGGACGCAACT
Zebrafinch    GCGCACGCCAGAGTTTCCTCTGCGGAACGGGCACGGCT
              * *** *     ******* *  **** ** * *
```

**Figure S3: Alignment of conserved enhancer-like sequence in the *IL10* locus of different avian species.** A 6kb genomic DNA sequence upstream of the first exon of the chicken *IL10* locus (Position Chr 26;2,564,995, GRCg6a:CM000117.5) was used to identify conserved sequences in multiple avian genomes using BLAST. A single highly conserved region was located around 2.3-2.8 kb upstream of the *IL10* transcription start site in each species shown. The sequences were aligned using ClustalW. Candidate purine-rich binding sites for ETS/NFAT family transcription factors (red), AP1 (turquoise box) and FOXP3 (pink box) are indicated.

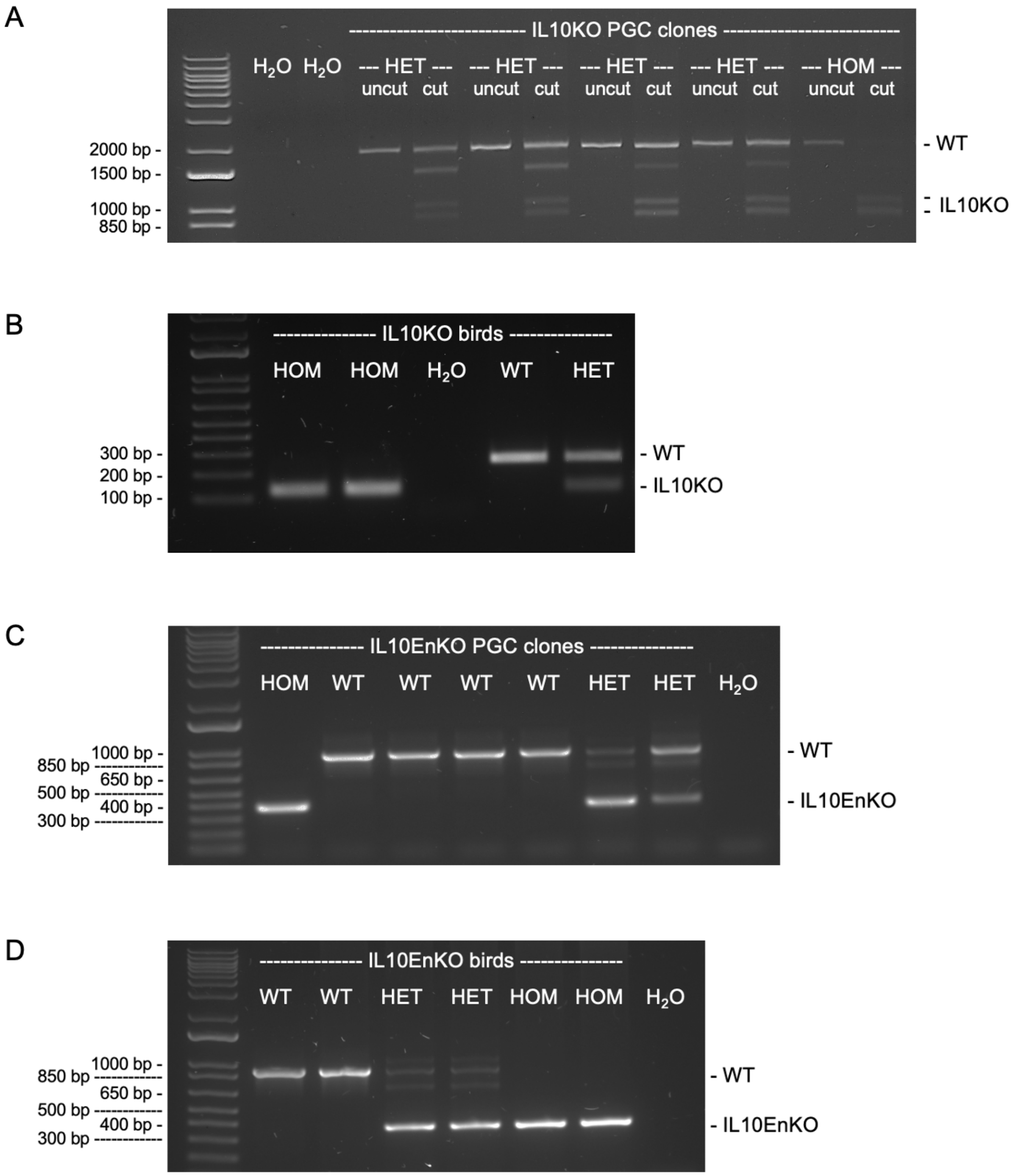


**Figure S4: Genotyping strategy for IL10KO and IL10EnKO edited PGCs and birds. A**: Primers IL10_Exon1_F4 and IL10_Exon1_R4 were used to amplify a 1826-bp fragment encompassing *IL10* exon 1 (uncut), followed by digestion with AvrII (cut) to identify PGC clones carrying the IL10KO edited allele. **B**: Primers IL10_Exon1_F1 and IL10_Exon1_R1 were used to amplify a 246-bp fragment encompassing *IL10* exon 1, followed by digestion with AvrII to identify birds carrying the IL10KO edited allele (gel shows PCR products after AvrII digestion). **C**: Primers IL10-Enhancer_F2 and IL10_Enhancer_R2 were used to amplify the *IL10* putative enhancer region and identify PGC clones carrying the IL10EnKO edited allele. **D**: Primers IL10-Enhancer_F2 and IL10_Enhancer_R2 were also used to identify birds carrying the IL10EnKO edited allele. See SI Appendix, Table S3 for primer sequences and expected fragment sizes.

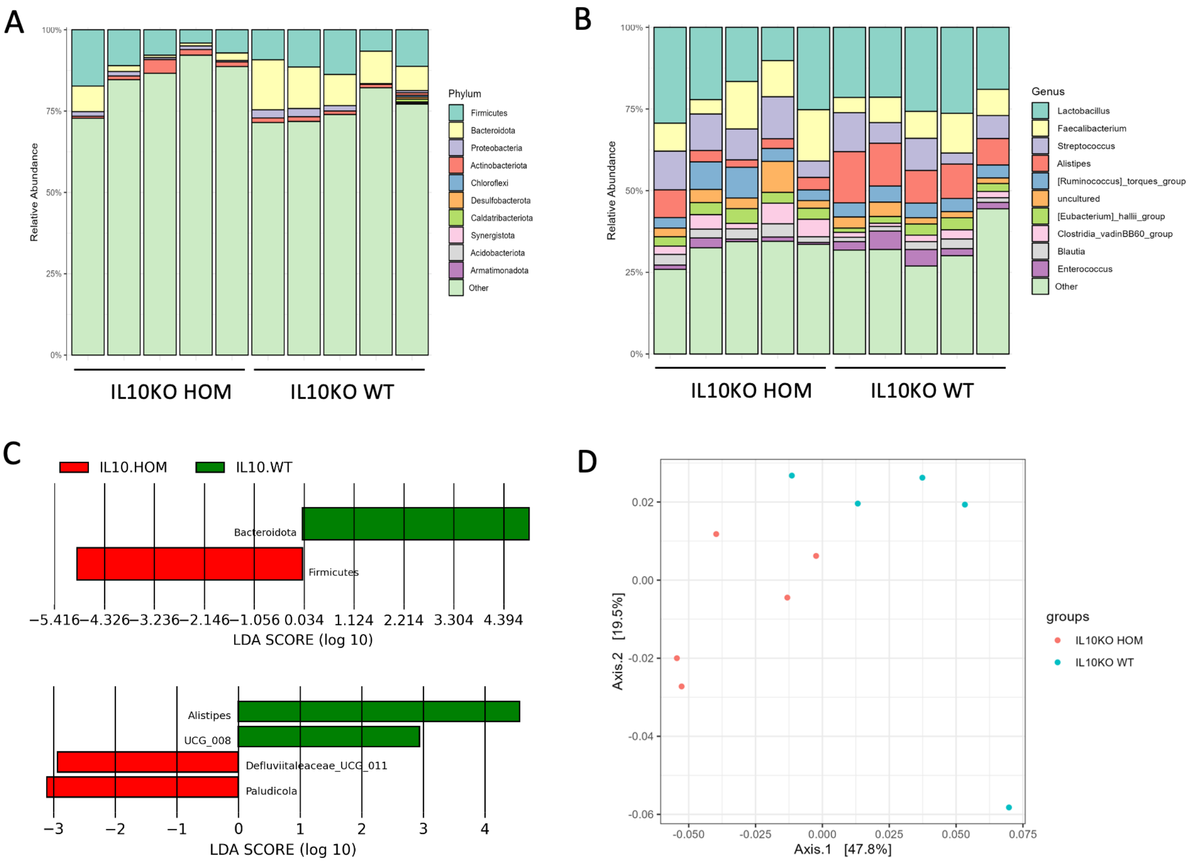


**Figure S5: Impact of IL10KO HOM mutation on the caecal microbiota of chickens.** Sequencing of 16S rDNA variable regions amplified from DNA extracted from the caecal contents of five birds of each genotype at four weeks of age revealed differences in the relative abundance of microbial phyla (**A**) and genera (**B**). Each vertical bar shows data from a different bird. LefSe analysis identified differentially abundant taxa (**C**). Beta diversity by principal component analysis based on the weighted UniFrac distance matrix (**D**).

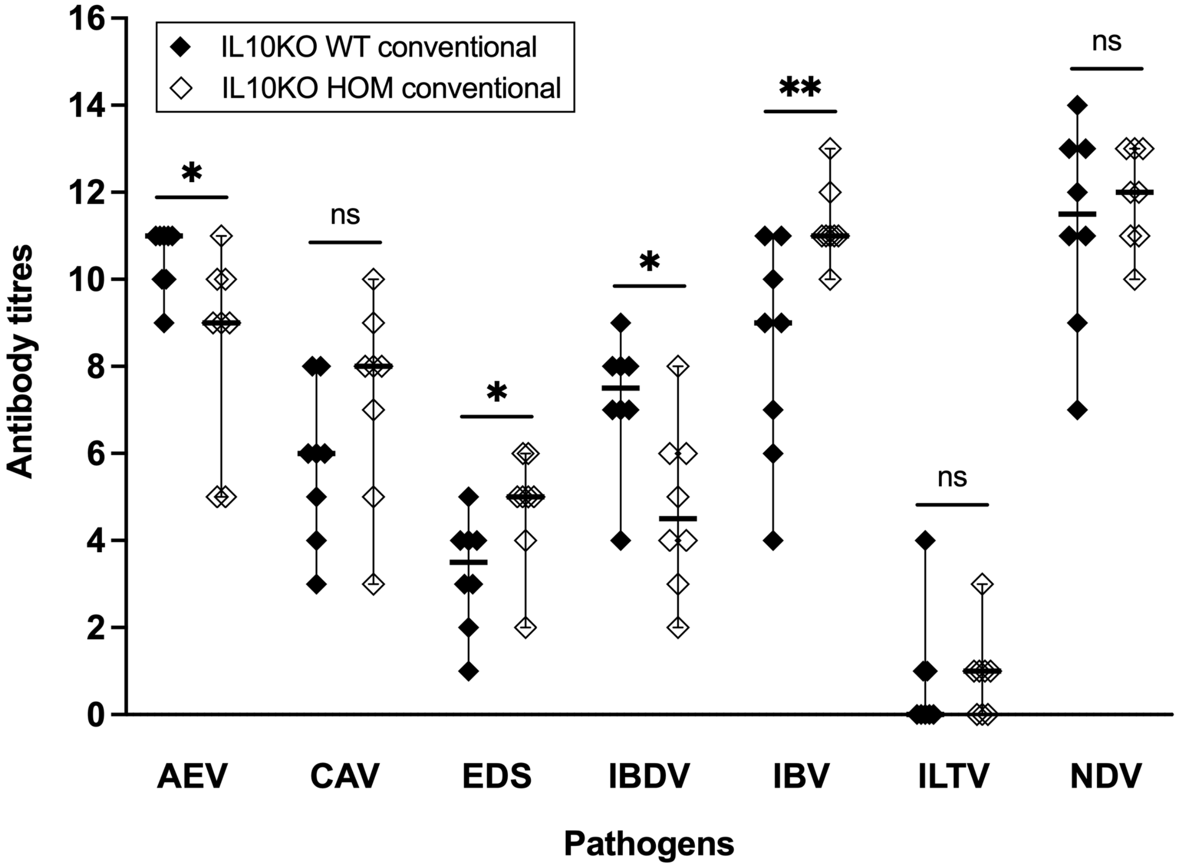


**Figure S6: Response to vaccination in IL10KO WT and HOM chickens.** Blood samples were collected from 29 weeks-old IL10KO WT and HOM vaccinated hens raised in the NARF conventional facility (n=8 in each group) and antibody titres were measured by ELISA. Titres to AEV and IBDV were significantly lower in IL10KO HOM hens compared to WT controls, whereas titres to EDS and IBV were significantly higher in IL10KO HOM hens compared to WT controls; titres to CAV, ILTV and NDV were not significantly different between IL10KO HOM and WT hens. Data displayed as median with 95% confidence interval. Statistical significance calculated using Mann-Whitney U tests; $*P < 0.05$, $**P < 0.01$, ns: not significant.

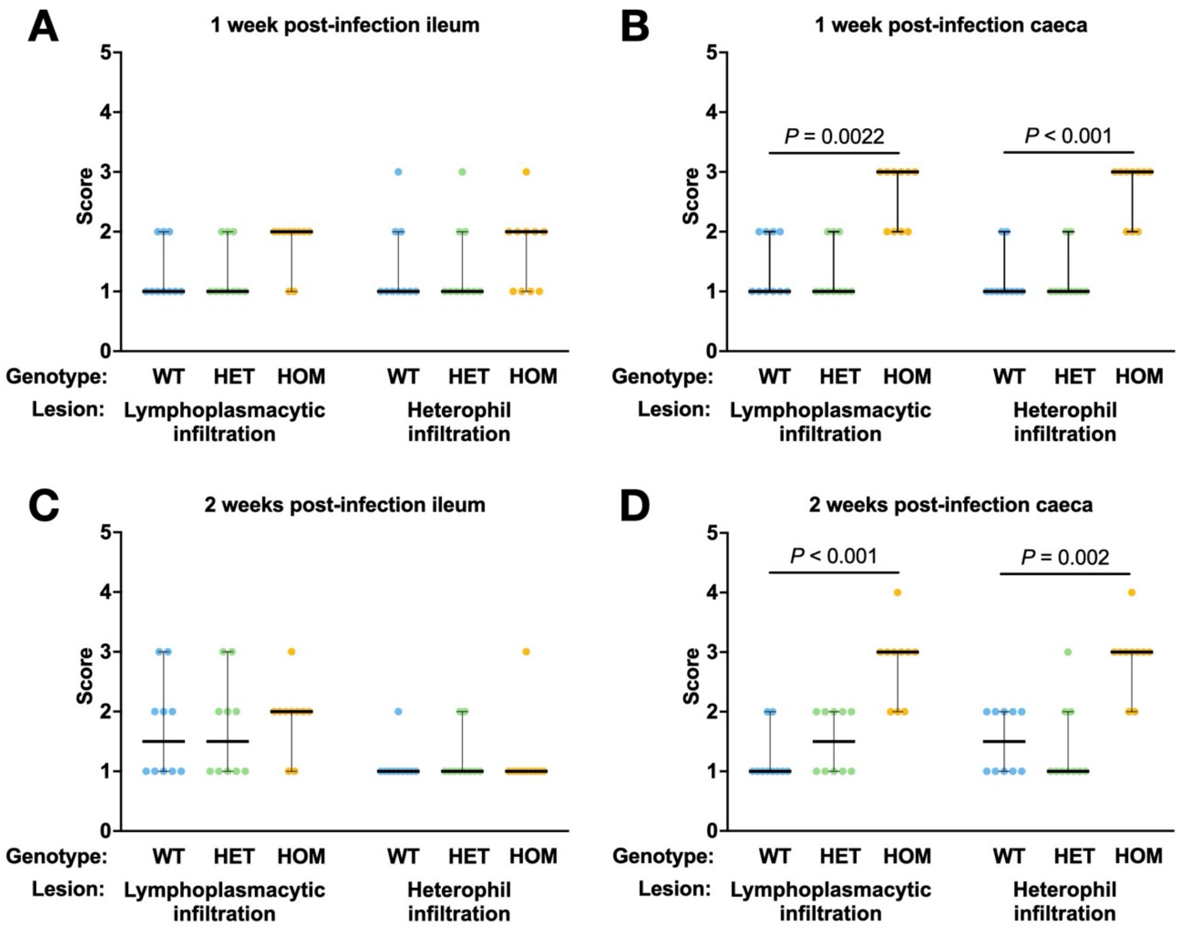


**Figure S7: Histopathological findings in ileum and caecum of IL10KO WT, HET and HOM chickens infected with *C. jejuni* cohort 1.** Histological sections were scored blind according to the scoring system described in Materials & Methods, by evaluating the extent of lymphocyte, plasma cell and heterophil infiltration. *P* values are shown where statistical differences between groups were detected.

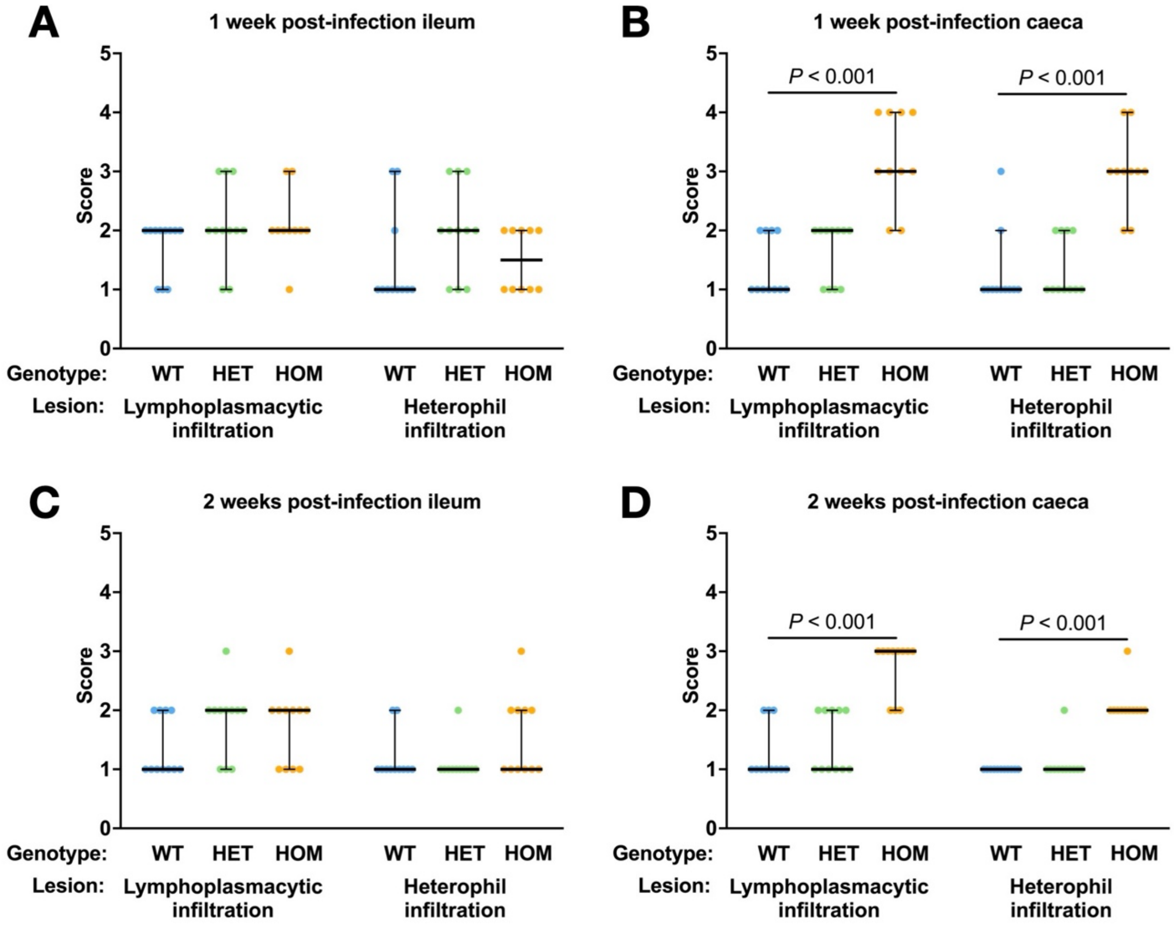


**Figure S8: Histopathological findings in ileum and caecum of IL10KO WT, HET and HOM chickens infected with *C. jejuni* cohort 2.** Histological lesions were scored in an identical fashion to cohort 1.

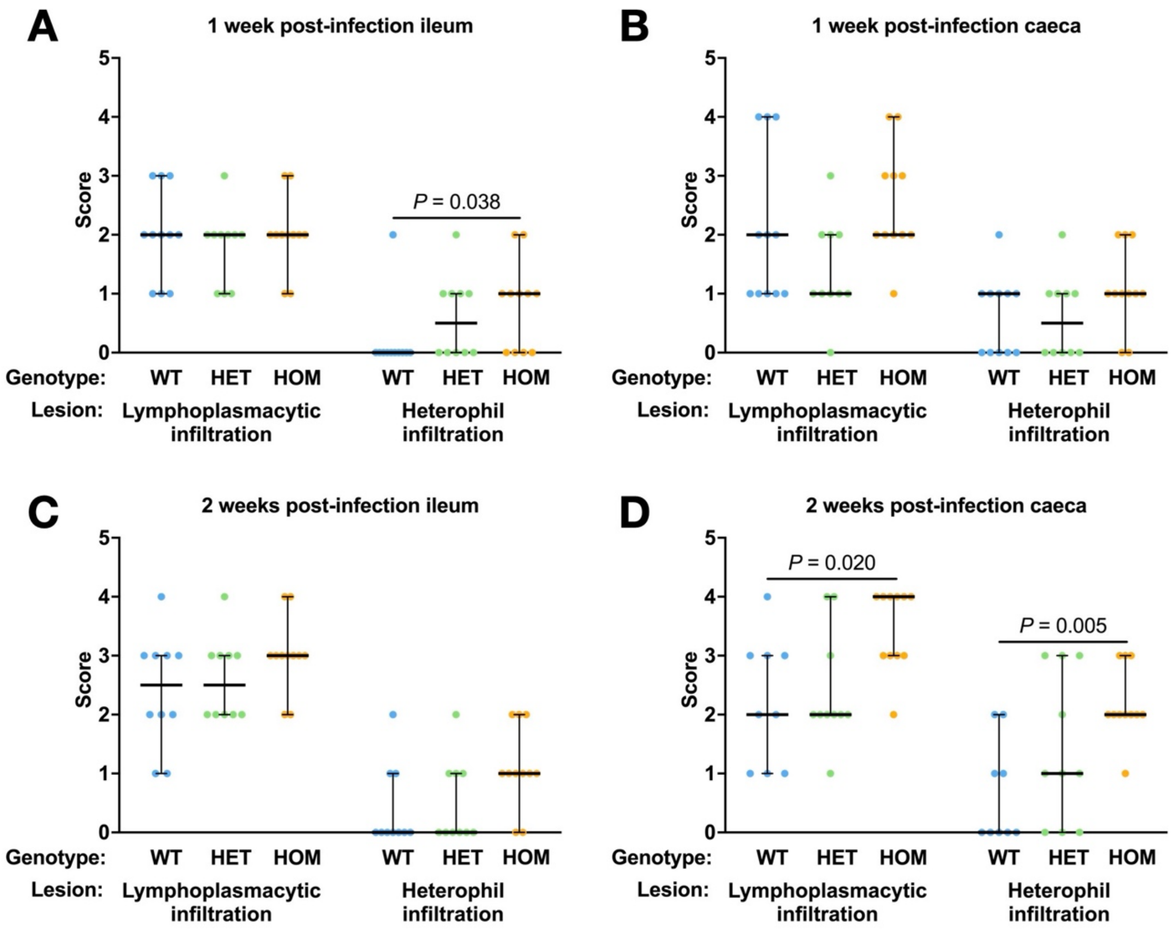


**Figure S9: Histopathological findings in ileum and caecum of IL10KO WT, HET and HOM chickens infected with *S.* Typhimurium.** Histological lesions were scored in an identical fashion to *Campylobacter* infection.

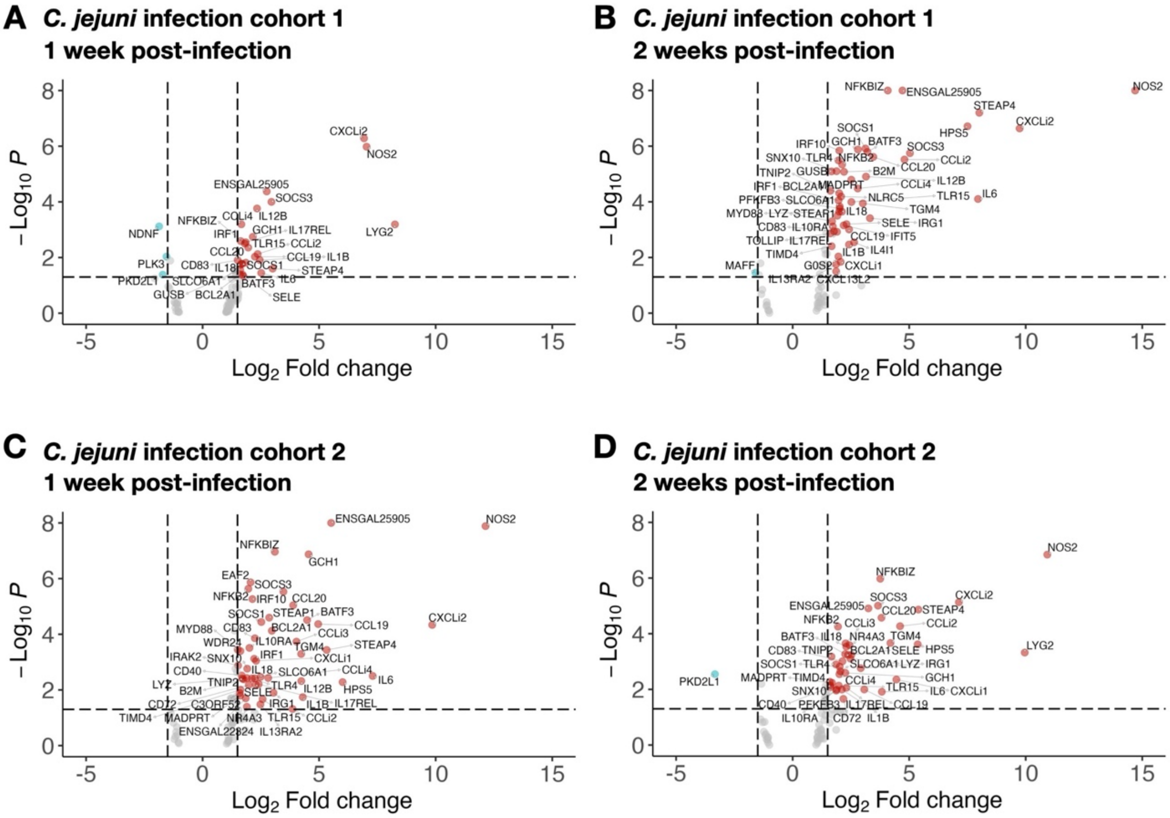


**Figure S10: Differential gene expression in caecum of IL10KO WT and HOM chickens infected with *C. jejuni*.** Volcano plots showing the direction and magnitude of changes in the transcription of immunity-related genes in the caeca of IL10KO WT versus IL10KO HOM chickens following infection with *C. jejuni* in cohort 1 (A, B) and cohort 2 (C, D).

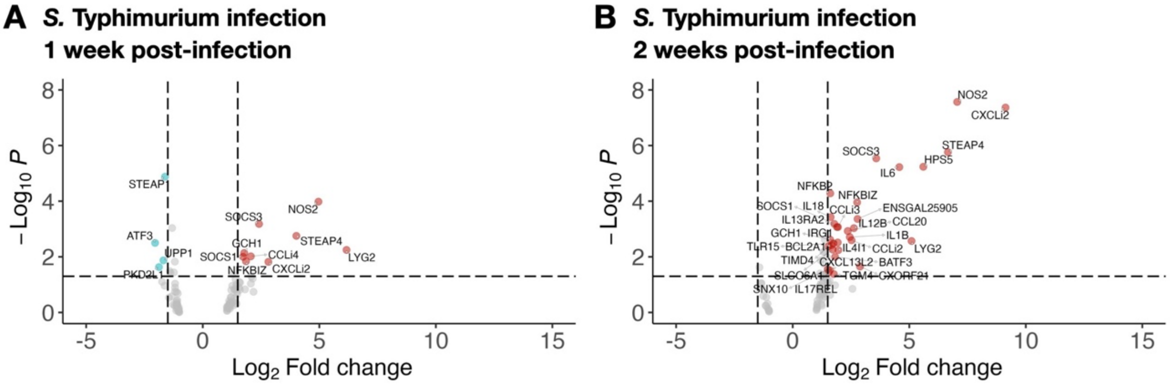


**Figure S11: Differential gene expression in caecum of IL10KO WT and HOM chickens infected with *S.* Typhimurium.** Analysis of differential transcription was identical to that of *Campylobacter* infection.

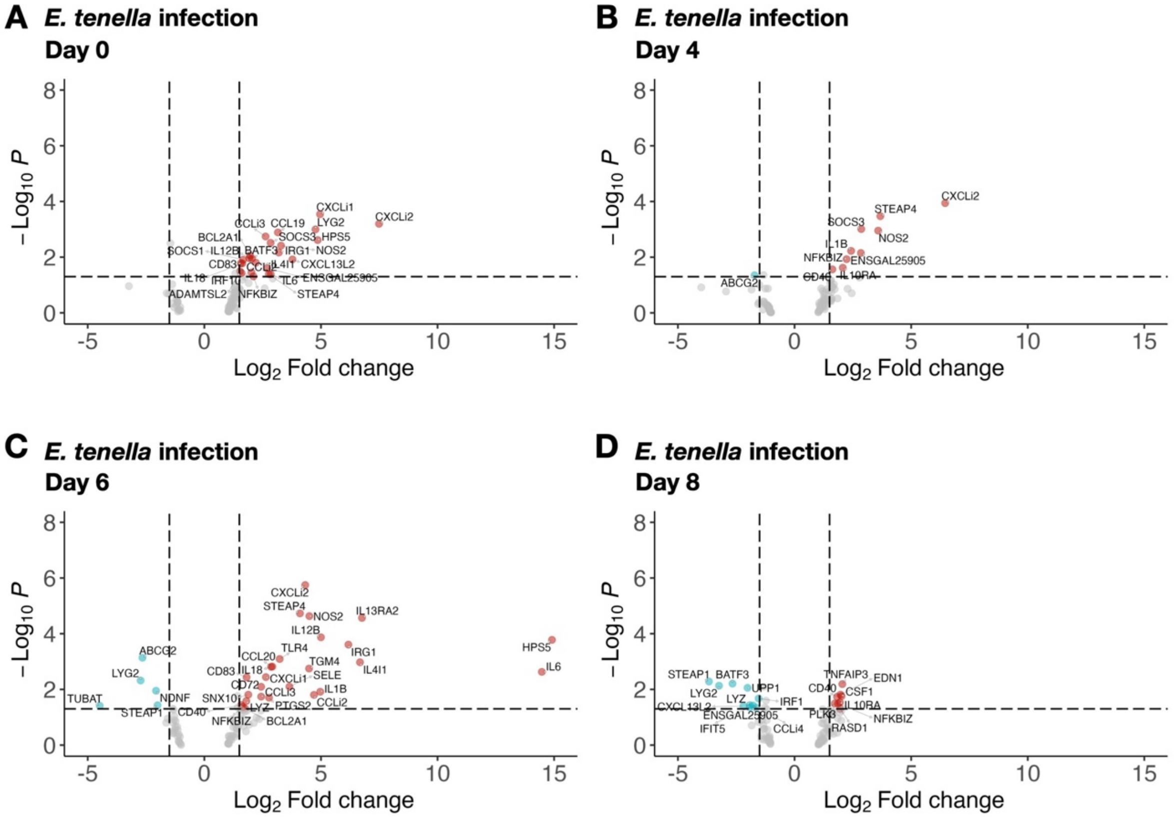


**Figure S12: Differential gene expression in caecum of IL10KO WT and HOM chickens infected with *E. tenella*.** Changes in transcription were analysed on days 0, 4, 6, and 8 post-infection.

**Table S1**: Number of IL10KO and IL10EnKO WT, HET and HOM chicks hatched in the NARF SPF chicken facility in the first (G1) and second (G2) generations. n.d.: not determined.

| Line | Parental cross | Generation | Total chicks hatched | Number of chicks per genotype | | | |
|---|---|---|---|---|---|---|---|
| | | | | WT | HET | HOM | n.d. |
| IL10KO | iCaspase9 surrogate host (G0) x RIR | G1 | 102 | 0 (0%) | 102 (100%) | 0 (0%) | 0 (0%) |
| | HET (G1) x HET (G1) | G2 | 435 | 105 (24%) | 209 (48%) | 94 (22%) | 27 (6%) |
| IL10EnKO | iCaspase9 surrogate host (G0) x RIR | G1 | 102 | 0 (0%) | 102 (100%) | 0 (0%) | 0 (0%) |
| | HET (G1) x HET (G1) | G2 | 246 | 62 (25%) | 104 (42%) | 67 (27%) | 13 (5%) |

**Table S2**: Routine vaccination schedule in the NARF conventional chicken facility.

| Age at vaccination | Vaccines | Type of vaccine | Route of administration | Pathogens |
|---|---|---|---|---|
| Day old | Prevexxion RN | Live | Intra-muscular injection | Marek's disease virus (MDV) |
| 5-7 days | Evant | Live | Drinking water | Avian coccidiosis virus (ACV) |
| 3-5 weeks | Nobilis Gumboro D78 | Live | Drinking water | Infectious bursal disease virus (IBDV) |
| 3-5 weeks | Poulvac IB Primer | Live | Drinking water | Infectious bronchitis virus (IBV) |
| 3-5 weeks | Nobilis ND Clone 30 | Live | Drinking water | Newcastle disease virus (NDV) |
| 6-9 weeks | Poulvac ILT | Live | Eye drop | Infectious laryngotracheitis virus (ILTV) |
| 6-12 weeks | Poulvac AE | Live | Drinking water | Avian encephalomyelitis virus (AEV) |
| 8-12 weeks | AviPro Thymovac | Live | Drinking water | Chicken anaemia virus (CAV) |
| 9-14 weeks | Nobilis IB Ma5 | Live | Drinking water | Infectious bronchitis virus (IBV) |
| | Nobilis IB 4-91 | Live | Drinking water | |
| 9-14 weeks | Nobilis ND Clone 30 | Live | Drinking water | Newcastle disease virus (NDV) |
| 16 weeks and over | Nobilis RT+IBmulti+ND +EDS | Inactivated | Intra-muscular injection | Avian rhinotracheitis virus (ARV) |
| | | | | Infectious bronchitis virus (IBV) |
| | | | | Newcastle disease virus (NDV) |
| | | | | Duck adenovirus, the agent of egg drop syndrome (EDS) |

**Table S3**: PCR primer sequences and expected fragment sizes.

| Primer name | Sequence 5'-3' | Expected fragment sizes | |
|---|---|---|---|
| | | WT allele | Edited or transgenic allele |
| Myco3_14709 | GCG GTG TGT ACA AGA CCC GA | ~500 bp (MS and MG) | N/A |
| Myco5_14712 | TGC CTG AGT AGT ACA TTC GC | | |
| Myco5_14713A | CGC CTG AGT AGT ATG CTC GC | | |
| IL10_Exon1_F1 | TAACCCCACGAAACAGAAGGAG | 243 bp | ~115 and 128 bp (after AvrII digestion) |
| IL10_Exon1_R1 | ATCAAGGGCAGCAGCAGAATAA | | |
| IL10_Exon1_F4 | AAGATGTGCTTTATGCAGGCAG | 1826 bp | ~855 and 965 bp (after AvrII digestion) |
| IL10_Exon1_R4 | GGGAAGCATGCAGAACTGAC | | |
| IL10-Enhancer_F2 | GGGTCGTCCCACACTTTCCT | 912 bp | ~380 bp |
| IL10-Enhancer_R2 | GAGTCAAGCTGCAAATTCCGAC | | |
| LTR_U3_F | TCCTCTGGTTTCCCTTTCGC | 320 bp | 400 bp |
| 235_F | CAGCCCACCCATCTCATCTC | | |
| 235_R | GACATTTCCTGCCTCCCCAG | | |

**Table S4**: Guide RNA and ssODN sequences. Note the three substituted nucleotides (red, lowercase) and AvrII restriction site (CCTAGG, underlined) in the ssODN sequence.

| Target | Name | Sequence 5'-3' |
|---|---|---|
| *IL10* exon 1 | ggIL10_exon1_g3 | GCAGCAGCTCAGAGAAGTGC |
| | IL10_exon1_HDRoligo2 | ACCTGCTGCCAAGCCCTGTTGCTGCTGCTGGCTGCATGCACCCTGCCTGCCCACTGCTTGGAGCCCACCTaggTGCACTTCTCTGAGCTGCTGCCCGCCCGGCTGCGGGAGCTGAGGGTGAAGTTTGAGGAAATTAAGGACTA |
| *IL10* putative enhancer region | IL10_enhancer_guide1 | GTTTCGTAGCGGGTGAATGA |
| | IL10_enhancer_guide2 | GTGCAGGGCAGTTTCCTTTG |

**Table S5**: NARF SPF screening.

| Pathogens | ELISA | PCR |
|---|---|---|
| 1. Avian adenovirus (FAV-1) | X | |
| 2. Avian encephalomyelitis virus (AEV) | X | |
| 3. Avian influenza virus (AIV) | X | |
| 4. Avian leukosis virus (ALV) | X | |
| 5. Avian pneumovirus (APV) | X | |
| 6. Avian reovirus (REO) | X | |
| 7. Avian reticuloendotheliosis virus (REV) | X | |
| 8. Chicken anaemia virus (CAV) | X | |
| 9. Duck adenovirus, the agent of egg drop syndrome (EDS) | X | |
| 10. Infectious bronchitis virus (IBV) | X | |
| 11. Infectious bursal disease virus (IBDV) | X | |
| 12. Infectious laryngotracheitis virus (ILTV) | X | |
| 13. Marek's disease virus (MDV) | | X |
| 14. Mycoplasma gallisepticum (MG) | X | |
| 15. Mycoplasma synoviae (MS) | X | |
| 16. Newcastle disease virus (NDV) | X | |
| 17. Salmonella gallinarum & pullorum | X | |
